# Transcriptomic profile of a rat jaw opener (anterior digastric) and a jaw closer (superficial masseter)

**DOI:** 10.64898/2026.08.28.747950

**Authors:** Prabath S. Meemaduma, Brandon P. Reder, Nicolai Konow, Jeffrey R. Moore, Matthew J. Gage

## Abstract

Mammalian skeletal muscle research predominantly focuses on locomotor muscles, and feeding related muscles remain less extensively characterized despite their role in mastication, mandibular stabilization, and swallowing. In this study, we investigated the transcriptomic specialization of three functionally and developmentally unique rat muscles: the anterior digastric (AD), a jaw opening muscle; the superficial masseter (SM), a jaw closing muscle; and the Sternohyoid (SH), a non-mandibular muscle involved in swallowing. Differential gene expression and weighted gene co-expression network analysis were used to characterize the transcription level features associated with their distinct roles. Our results indicated that all three muscles predominantly expressed fast-twitch contractile isoforms. However, the AD showed lower overall expression of several contractile gene families, including myosin heavy chain, myosin light chain, and tropomyosin isoforms, while exhibiting elevated expression of slow/oxidative myosin isoforms like *Myh7* and *Myh2*. Network analysis revealed that modules correlated with AD are strongly enriched for fatty acid catabolism, mitochondrial energy production, and vascular/extracellular matrix remodeling. Additionally, AD and SM shared a distinct gene set compared to SH, highlighting their common developmental origin from the first branchial arch. Our findings show that the rat feeding related muscles possess unique transcriptomic profiles shaped by their contractile functions, developmental origins, and metabolic functions.

## Introduction

Mammalian skeletal muscle studies can be broadly categorized into two groups: feeding related muscles and locomotor muscles. Many muscle related studies have focused on locomotion due in part to the number of diseases connected to deficiencies of locomotor muscle function. In contrast, feeding muscles has received less attention despite their importance in biting, swallowing, and mastication. Retaining masticatory and swallowing functions become particularly important with age because individuals must continue to eat and swallow even after their locomotor capacity has substantially declined. In a steadily aging population, dysphagia is becoming an increasingly prevalent clinical and public health challenge that can compromise the nutritional status of the elderly population [19]. While the fundamental principles governing muscle contraction are maintained between locomotion and feeding, comparisons between the two families of muscles have revealed that jaw muscles show faster contractile velocities and faster twitch times [1]. Other studies on mammals have proposed that jaw muscles are adapted to precise and high forces at the expense of excursion and velocity compared to efficiency focused movement in locomotor muscles [10]. These differences, while not surprising due to their different physiological roles, highlight the importance of understanding the underlying transcriptomic makeup of these muscle types and how these may differ from locomotor muscles.

The model system used in this study is *Rattus norvegicus* (Rat) due to the significant body of existing research exploring the prime jaw opener - anterior digastric (AD) and one of jaw closer muscles superficial masseter (SM) [12, 22, 23]. The anterior digastric is a muscle responsible for depression of the mandible during jaw opening and also contributes to hyoid movement. The superficial masseter functions antagonistically to AD in closing the jaw by elevating the mandible. Both the AD and the SM are derived from the first branchial arch. Sternohyoid (SH) which moves and stabilizes the hyoid but is not directly involved in mastication. SH was included as both a functional and developmental control because it is not derived from the first branchial arch.

Skeletal muscle fibers are categorized into two major categories based on their contractile speeds and length of contractions. Type I fibers have slower twitch/contractile speeds but are fatigue-resistant due to their oxidative respiration. Type II fibers have faster twitch speeds and fatigue faster than slow twitch fibers due to their glycolytic respiration [14]. Fast-twitch fibers would often favor fast myosin heavy isoforms MyHC-2X, MyHC-2A and MyHC-2B. These proteins are encoded for by the *Myh1, Myh2*, and *Myh4* genes, respectively. The slow-twitch myosin heavy chain isoform (MyHC-*β*) is encoded by gene *Myh7*. Rat jaw muscles mainly express fast-type fibers. The SM muscles are known to exclusively express fast-twitch type II myosin heavy chain (MHC) isoforms (primarily IIx, IIa, and IIb) and completely lack slow twitch type I fibers [3]. In contrast, AD muscles express a more diverse profile consisting of low levels of type I, along with fast-type IIa, IIb, and IIx isoforms [12].

Compared to jaw closing superficial masseter muscles, the jaw-opening anterior digastric exhibits higher calcium sensitivity and energy efficiency, properties attributed to its elevated expression of slow-type fibers [23]. These physiological characteristics correlate with the daily muscle activity (duty time) patterns observed in rats: the digastric muscles have a longer duty time and lower intensity contractions than masseter which shows lower duty time at higher intensities [23]. The fatigue resistance of AD is well suited to the prolonged, low-intensity activity required for functions such as postural stabilization of the mandible. In contrast, the rat jaw-closers such as superficial masseter perform the high intensity, phasic contractions required to bite down on food and therefore contain a greater proportion of fast-type fibers [12]. The underlying transcriptomic landscape that regulates this specialization and low abundance of regulatory molecules such as transcription factors and signaling molecules remain unexplored.

## Materials and Methods

### Sample acquisition and preparation

Sprague Dawley rats sourced from Charles River. Same-sex pairs were housed under a 12H light:12H dark cycle at the Harvard University Concord Field Station. All methods are in accordance with ARRIVE guidelines. Animal use and experimental methods were approved by the Institutional Animal Care and Use Committees at Harvard University (FAS, 20-09 04) and UMass Lowell (23-03-KON). The rats were kept on water and rodent chow *ad libitum*, and the muscle tissue were harvested between 11 and 16 weeks of age. Rats (n=9) were euthanized by isoflurane overdose followed by heart removal. The jaw muscle tissue was dissected and surgically cut into pea sized portions and immediately transferred into a vial containing RNALater solution and stored at −20° C prior to RNA extraction.

### RNA extraction, cDNA library preparation and RNA sequencing

Total RNA was extracted using a RNeasy Fibrous Tissue Mini Kit (QIAGEN, 74704) and quantified using a Qubit RNA Broad-Range assay. The fragment size profile was analyzed using an Agilent 4200 TapeStation System. Extracted RNA samples with an RNA integrity value (RIN) of 7 or above were used in subsequent library preparation. Input material was normalized to 250 ng, and the cDNA library generation was carried out using Illumina TruSeq Stranded mRNA Kit with IDT for Illumina TruSeq UD Indexes. Sequencing was carried out on an Illumina NextSeq 550 sequencer using 150 paired end cycles.

### Data preprocessing and alignment

The raw paired-end RNA-seq reads were filtered and trimmed using Trimmomatic v. 0.39 [2]. The adapter sequences were removed with ILLUMINACLIP using the adapter file TruSeq3-PE.fa. The maximum seed mismatch count was set at 2, and palindrome clip threshold was set at 30, and simple clip threshold was set at 10.

The 3 leading and 3 trailing low-quality bases from the beginning and end were also removed. Following this step, a sliding window trimming was conducted using a window size of 4 and a quality threshold of 15. Reads with lengths shorter than 100 bases were also discarded.

The trimmed and filtered paired-end reads were aligned to the *Rattus norvegicus* RefSeq GRCr8 genome assembly (GCF 036323735.1) using STAR aligner v. 2.7.10b [6]. The BAM output from STAR was used in calculating the Transcripts per kilobase million (TPM) values using RSEM [17]. FastQC (https://www.bioinformatics.babraham.ac.uk/projects/fastqc/) was used to assess the sample read quality, adapter trimming, and mapping percentages. Samples with fewer than 13 million mapped reads were excluded in all subsequent analysis (Average library size = 19.04M, range: 13.94M-26.98M). The final dataset consisted of 8 AD, 6 SH, and 6 SM replicates.

### Data analysis and differential gene expression analysis

The raw gene counts generated by STAR were imported using the readDGE function of the edgeR (v4.2.2) library [4]. To minimize low-abundance background noise, genes with minimal expression were filtered out across samples using the filterByExpr function with specified minimum count of 10 and minimum proportion of samples set at 0.7. This filtering pipeline reduced the initial dataset from 38,673 down to 12,962 genes. Genomic annotations including Entrez gene IDs, Ensembl gene IDs, and gene descriptions were appended to the filtered matrix using the biomaRt library [7].

Counts per million (CPM) values were calculated using edgeR. The limma (v3.60.6) library [24] was used both to correct for batch effects via removeBatchEffect and to visualize the CPM counts in a multidimensional scaling (MDS) plot generated using plotMDS. Differential expression testing was executed on the filtered raw count matrix using DESeq2 (v1.44.0) [18], incorporating the batch variable directly into the design formula. DESeq2 was also used to estimate sample size factors and normalize for sequencing depth. Log2 fold changes were determined between the tissue groups, and FDR was controlled using the Benjamini-Hochberg (BH) correction method. The resulting BH-adjusted p-values (padj) of these pairwise comparisons were rendered on the TPM barplots (∗*padj <* 0.05, ∗ ∗ *padj <* 0.01, ∗ ∗ ∗*padj <* 0.001, ∗ ∗ ∗ ∗ *padj <* 0.0001).

### Weighted gene co-expression network analysis (WGCNA)

Raw count files from STAR output were used in the construction of the network. The removeBatchEffect function of limma (v3.60.6) was used to remove the batch effect. Genes with low variance were filtered using varFilter in the genefilter (v1.86.0) package [8]. A 0.5 cutoff value was applied to exclude the bottom 50% of genes with the lowest expression variance. This value is used in several WGCNA studies to reduce background noise and improve network construction efficiency [20, 27]. This reduced the dataset from 12,962 down to 6,410. The samples were clustered using hclust function of fastcluster 1.3.0 and a sample tree was plotted from Euclidean distances to detect potential outliers [21]. A tree cut height of 90 was applied to exclude outlying samples in this dataset (Figure 1a). After filtering, there were 7 AD, 6 SH, and 7 SM samples (samples used in constructing the network are different from the samples used in the DE analysis).

**Figure 1.**
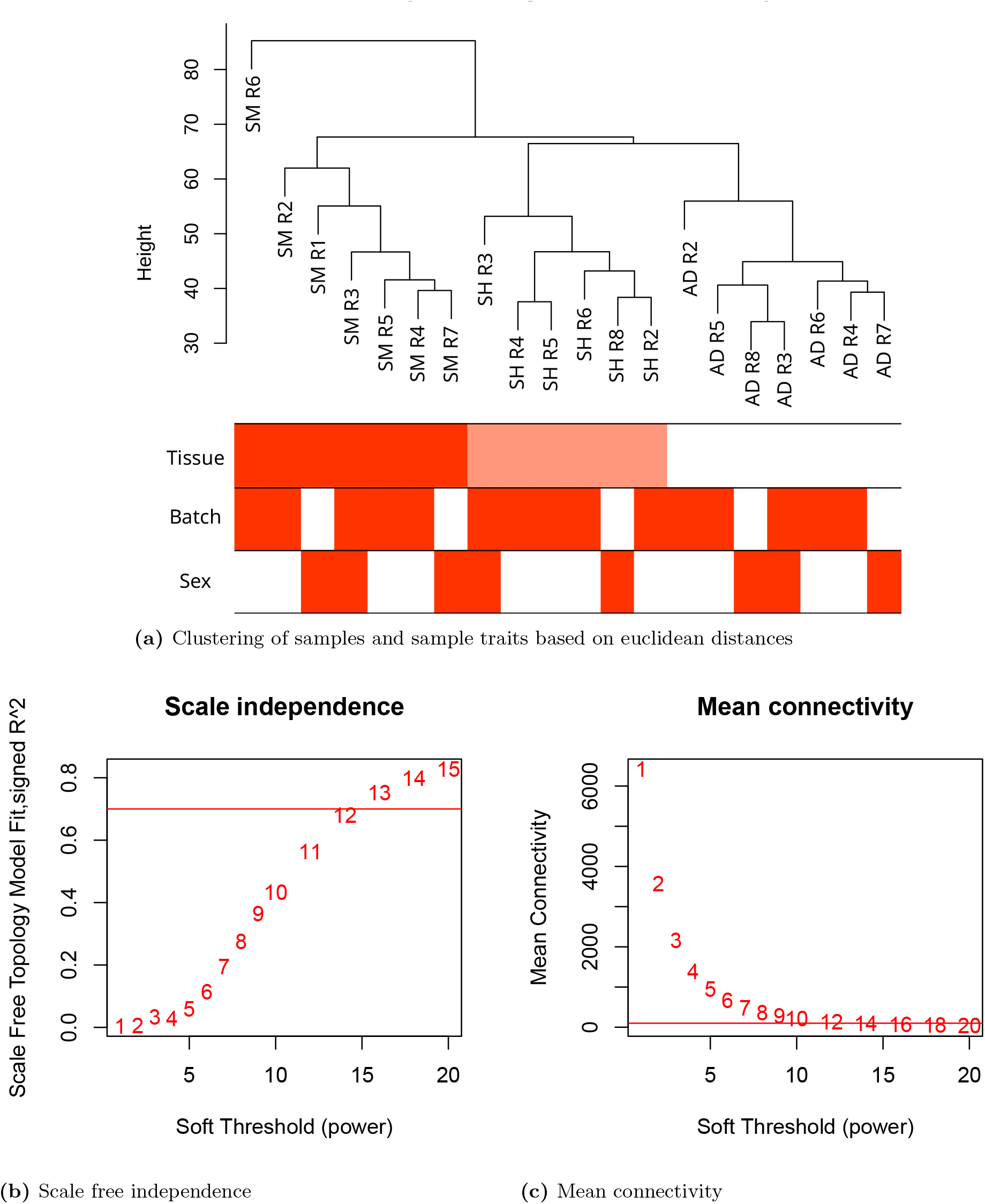
WGCNA sample tree, scale independence and mean connectivity

To construct the gene co-expression network, we utilized the WGCNA package in R [15]. We used the pickSoftThreshould function to test a range of values from 1-20 to select a soft thresholding power as shown in Figures 1b and 1c. Langfelder et al. [15] recommend selecting the first power between 1-20 achieving a scale-free topology R^2^ value of *>*0.7 and a mean connectivity between 100-200. Based on these criteria, a soft thresholding power of 12 was selected.

### Protein extraction, mass spectroscopy and database search

To extract proteins for the proteomics analysis, rat muscle tissues were homogenized and proteins were extracted using TRIzol™ Reagent according to the manufacturer’s protocol. In the final step, the protein pellet was resuspended in 23 µL of 1x S-trap lysis buffer, followed by protein reduction with 10 mM TCEP and alkylation with 20 mM iodoacetamide. The proteins underwent overnight digestion at 37 °C with trypsin. After digestion, 40 µL of 50 mM TEAB elution buffer was added and centrifuged at 4000 rpm for 1 minute, followed by a second 1 minute spin with 40 µL of 0.2% formic acid and a final spin with 50% acetonitrile. The resulting elution was dried using a speed-vac.

Liquid chromatography-tandem mass spectrometry (LC-MS/MS) was conducted using a TimsTOF Pro2 (Bruker) mass spectrometer coupled to a nanoElute LC system (Bruker) with 1 µL sample injections. For data processing, the custom ‘LFQ-MBR’ workflow from FragPipe (v21.1) was used. Database searches were conducted using MSFragger (v4.0, default parameters), with deep-learning prediction re-scoring executed via MSBooster (default parameters), Percolator (default parameters), and ProteinProphet (default parameters - Philosopher (v5.1.0)) for PSM validation and protein inference. The raw data files from Bruker were searched against the *Rattus norvegicus* (UP000002494) appended with common protein contaminants. Mass tolerances for precursor and fragment ions were set to 20 ppm, and the enzyme parameters were set to trypsin (not cutting before P) with an allowed peptide length of 7-30. Variable modifications were set to oxidation of Met and acetylation of protein N-t, while fixed modifications were set for carbamidomethylation of Cys. The results were filtered at a stringency of 1% False Discovery Rate (FDR) at the protein level. To identify the phosphorylated peptides, an additional search utilizing the Fragpipe/MSFragger trypsin specific workflow was conducted by including a phosphorylation modification (+79.9799) for the amino acids S, T, and Y.

For downstream analysis and quantification, the search outputs were imported into Scaffold Viewer (v5.3.3) to export the protein quantitative reports. The peptide entries in the report were then filtered by removing duplicate rows and the sum of Total Precursor Intensities (TPI) was calculated for each peptide across all samples. The top 5 peptides with the highest TPI were selected as the representative value for the abundances of each identified protein in the downstream data analysis. The final proteomic dataset only contained a single replicate for each tissue type and was utilized exclusively to corroborate the transcriptomic profiles.

## Results

A multi-dimensional scaling (MDS) plot was used to identify the main clustering patterns in the RNA-Seq dataset. Figure 2 shows the MDS plot using the top 10,000 genes by expression after batch correction. Each tissue type in the MDS plot forms distinct and non-overlapping clusters, and 95%-99% of sample variance lies within the ellipses (3 standard deviations). Samples were sourced from both female and male rats, as denoted by the ‘F’ and ‘M’ respectively on the MDS plot. Tissue type was identified as the primary driver of sample variance, since no distinct sex-based sub-clusters were observed within the ellipses. The first dimension of the MDS plot explains 23% of the variance in the dataset and separates SM samples from the AD and SH samples, thus separating the jaw closer SM from jaw opener muscles. Dimension 2 explains 17% of the variance and separates SH samples from the AD and SM samples, i.e., a pharyngeal arch separation.

**Figure 2.**
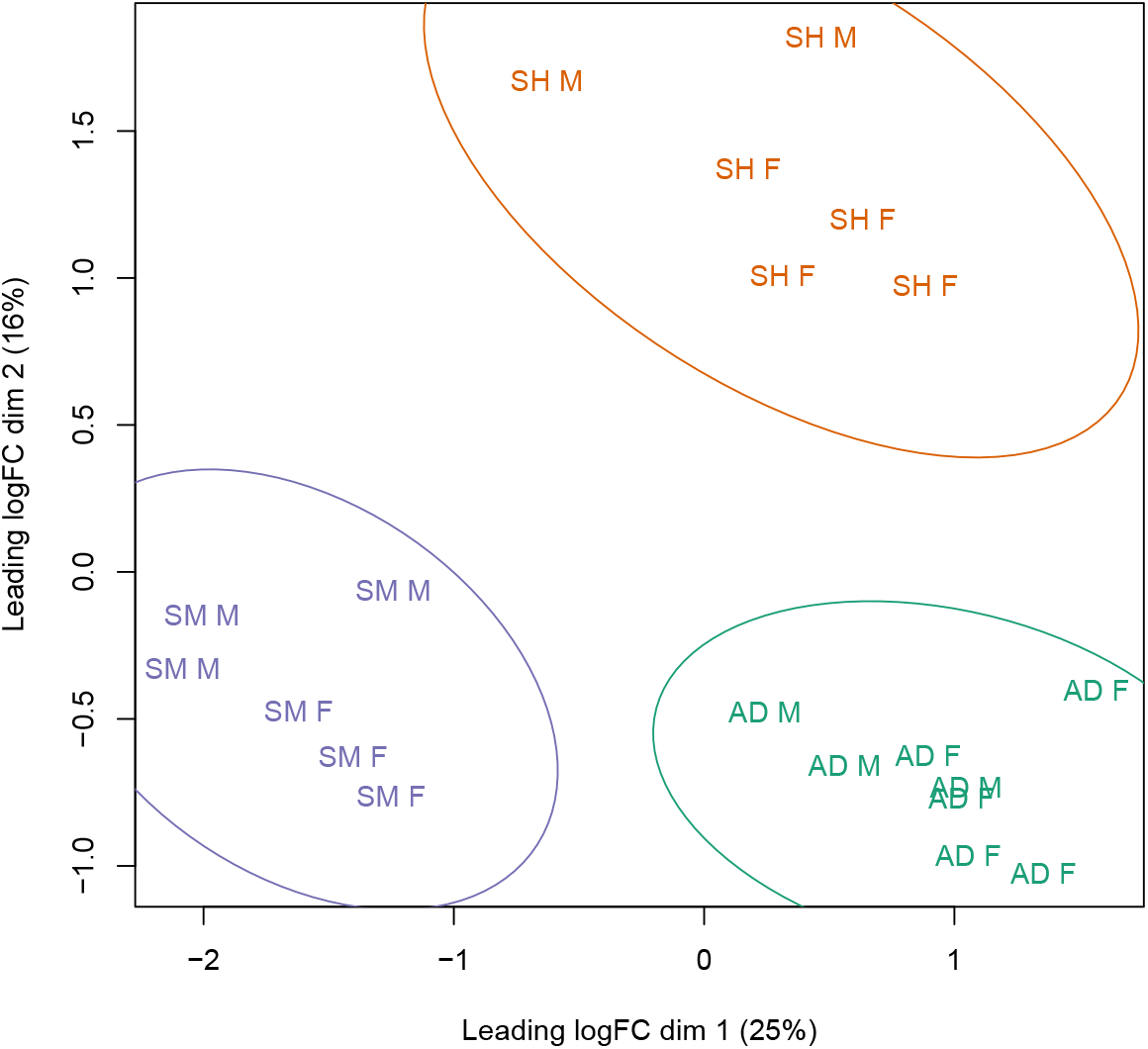
Leading logFC multidimensional scaling plot revealed three clusters separated by muscle tissue type AD (n = 8), SH (n = 6) and SM (n = 6). (Dimension1 = 25% variance and Dimension2 = 16% variance). Ellipses demarcate the 3 standard deviations of confidence boundaries calculated from the centroids of each tissue cluster.

### Rat anterior digastric expresses elevated levels of sarcomeric proteins compared to superficial masseter

Gene expression levels of the *Myh* isoforms are shown in Figure 3a in mean TPM ± SEM. The cumulative transcript counts of these four myosin isoforms are listed in Table 1. The myosin heavy chain expression is surprisingly similar (1.2% difference) between AD and SM. All three investigated muscle tissues preferentially express fast-twitch myosin heavy chain isoforms. Out of these fast-twitch isoforms, the glycolytic *Myh4* is the most abundant isoform, comprising over 50% of total Myh expression across all three muscles, followed by Myh1 (Table 2). Expression of the neonatal *Myh8* and other isoforms was negligible and was excluded from this list. The SM is exclusively characterized by the fast-glycolytic *Myh1* and *Myh4*. Predominant isoforms expressed in AD follow a similar baseline expression to SM muscles, the AD also exhibits significantly higher expression levels of *Myh2* compared to SM. Overall expression of the slow twitch *Myh7* isoform are low across the three muscles, the AD expresses higher proportions of both *Myh7* (oxidative) and *Myh2* (oxidative-glycolytic) compared to the other tissues studied. The 2% transcript level expression of *Myh7* in AD contrasts previous protein level expression studies in rat jaw muscles [23], which reported MHC-I levels closer to 10% in the AD.

**Table 1.** Sum of mean expression of contractile proteins in Transcripts per kilobase million. Overall transcript expression of AD is significantly lower in AD compared to SM.

|  | Gene isoform | AD (n=8) | SH (n=6) | SM (n=6) |
| --- | --- | --- | --- | --- |
| Total mean transcript counts in tpkm | *Myh* | 15,774 | 25,923 | 15,585 |
|  | *Myl* | 47,121 | 53,485 | 69,204 |
|  | *Tpm* | 29,182 | 34,593 | 40,476 |

**Table 2.**
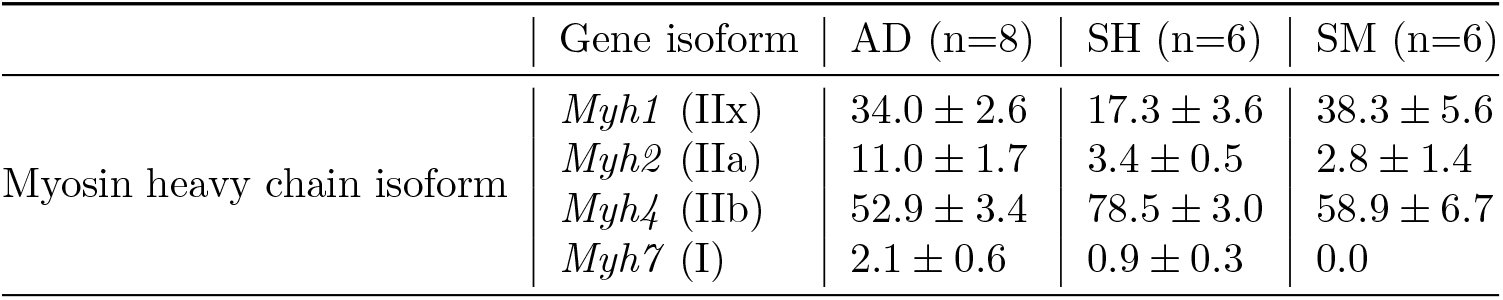
Myosin heavy chain isoforms percentages. AD, SH and SM predominantly express *Myh4* and *Myh1* while AD expression includes significantly higher levels of oxidative *Myh7* and oxidative-glycolytic *Myh2*.

**Figure 3.**
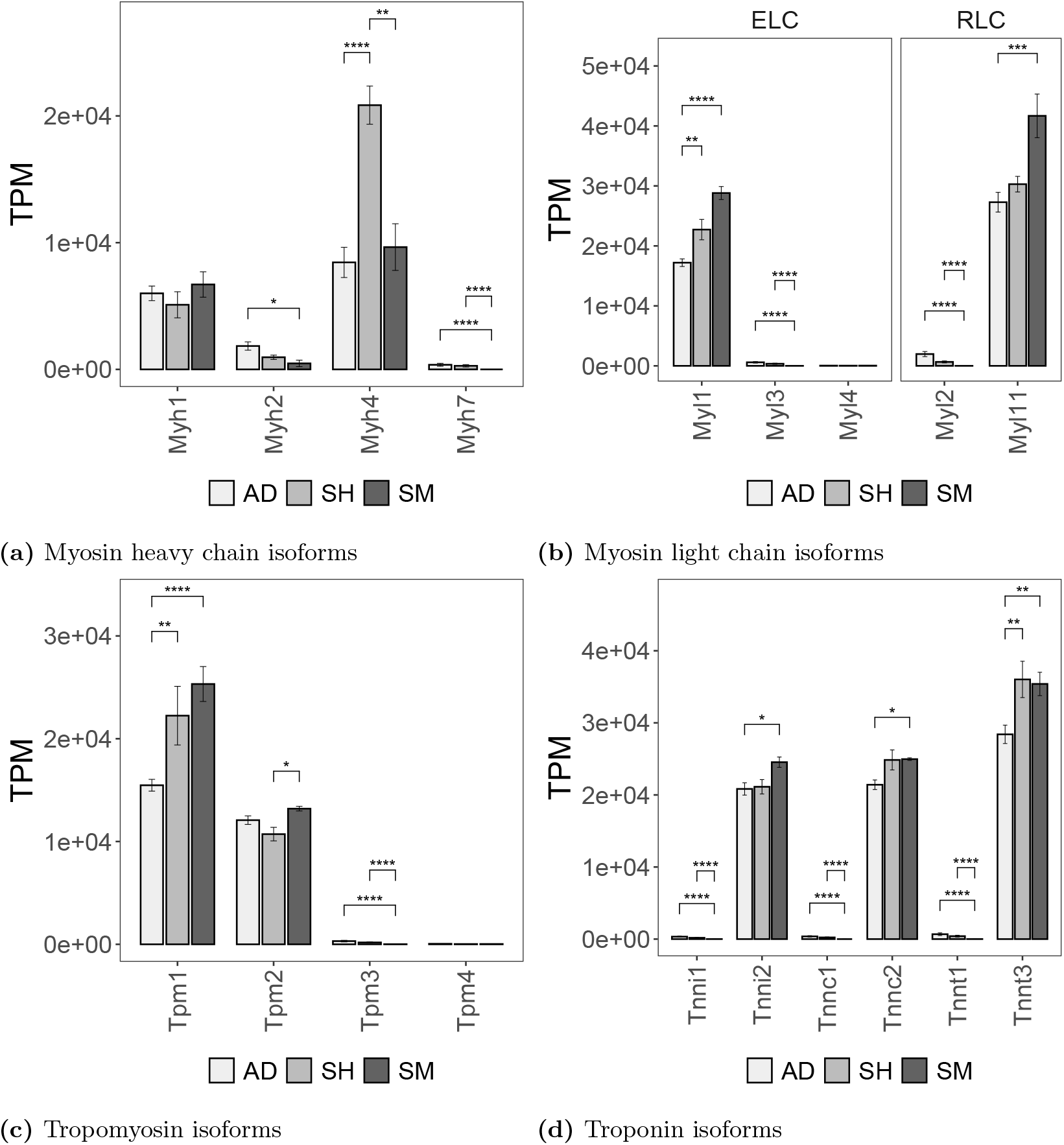
Transcriptomic expression profiling of major contractile protein isoforms across rodent muscles. Bar charts illustrate the transcript abundance quantified in Transcripts per kilobase million (TPM) and the three columns from left to right for each gene isoform is respectively anterior digastric (AD), sternohyoid (SH), and superficial masseter(SM). Brackets indicate the statistical significant differential expression (DESeq2 BH adjusted p ¡ 0.05). All three tissues predominantly express the fast isoforms of these contractile proteins.

The slow vs. fast fiber type characterization extends to other contractile proteins such as myosin light chain, tropomyosin, and troponin, where specific isoforms are predominantly expressed in each fiber type. *Myl1* is the fast isoform of essential light chain and *Myl11* is the fast regulatory light chain isoform. All three muscles primarily express the fast-twitch isoform of myosin light chains. AD muscles also express a low amount of *Myl2*, the slow type myosin light chain isoform. Similar to myosin heavy chain expression, cumulative myosin light chain expression is the lowest in AD at 47,121 TPM, whereas SM expresses a significantly higher level of *Myl* isoforms at 69,204 (log2FC = 0.55, p = 0.002). This expression pattern is observed in other major contractile proteins as seen in Figure 3 and Table 1. Fast skeletal isoforms (*Tnnc2, Tnni2*, and *Tnnt3* of all three subunits are dominant in all three muscles (Figure 3d). However, the expression profile of slow troponin isoforms (*Tnni1, Tnnc1*, and *Tnnt1*) is significantly upregulated in AD compared to the other two muscles.

### Differential expression analysis of Rat jaw opener shows heightened levels of calcium sensitivity and M-band genes

Differential expression analysis was carried out using DESeq2 to identify the genes with elevated or suppressed expression between SM and AD muscles. Figure 4 shows the design of these comparisons and the number of differentially expressed genes (DE genes) in each comparison. Initial filtering step using the filterByExpr function successfully removed genes that did not meet the minimum of 20 raw counts, in at least 6 samples (smallest group size) and genes that did not meet a minimum total count of 50 across all samples were filtered. The SM vs. AD comparison resulted in 132 genes up regulated in SM with a log2 fold change over 0.5 (padj ¡ 0.05) and 224 genes down regulated in SM compared to AD.

**Figure 4.**
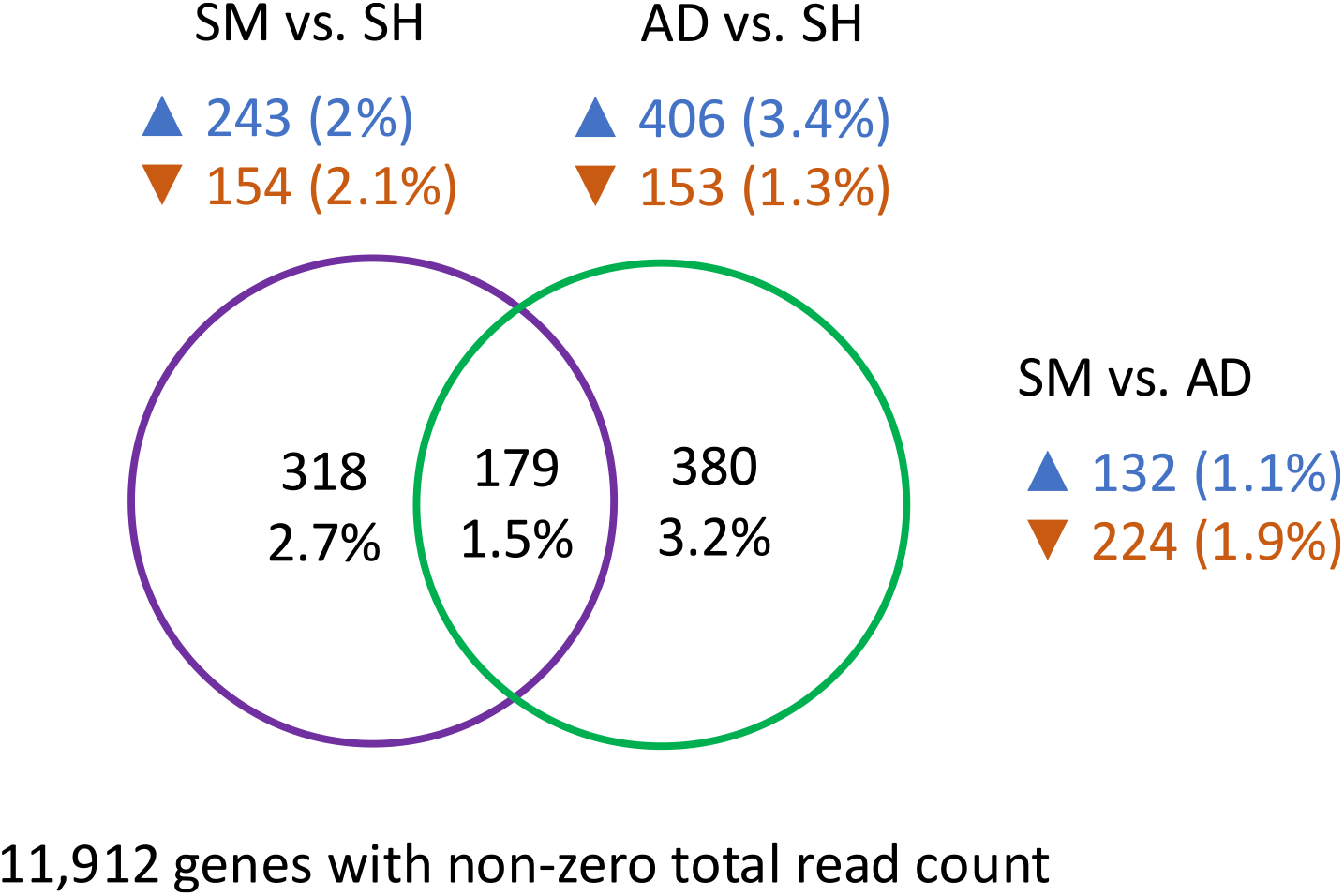
Pairwise differential expression profiling reveals AD and SM muscles exhibit fewer DE genes between each other than when compared to non-mandibular branchial arch SH. The figure shows upregulated and downregulated genes in the three comparisons: (A) SM vs. SH comparison showing number of genes upregulated in SM in blue and upregulated in SH in red; (B) AD vs. SH comparison showing number of upregulated genes in AD in blue and upregulated in SH in red; (C) SM vs. AD comparison, upregulated in SM in blue and upregulated in AD in red. The number of common genes expressed in comparisons involving SH are shown in the intersection areas of the two sets in the Venn diagram. These genes are commonly differentially expressed in both branchial arch muscles (AD and SM) compared to SH. The criteria for selecting significant gene expression was set at —log_2_FC— ¿ 0.5 and adjusted p-value ¡ 0.05

A conservative log2 fold change threshold of 0.5 was used since all three tissues are skeletal muscles with almost identical expression profiles as seen in previous results. The Venn diagram in Figure 4 shows the number of unique DE genes found in the comparisons between SM vs. SH and AD vs. SH, and the intersection showing the common DE genes between both comparisons. A total of 318 unique DE genes were identified in the SM vs SH comparison and 380 unique DE genes in the AD vs. SH comparison, with only 179 DE genes in common to both comparisons. It was predicted that the genes common to both these comparisons would highlight the unique expression of SH tissues compared to jaw muscle tissues derived from the first branchial arch.

AD vs. SM comparison yielded 369 DE genes and the enriched gene ontology terms (GO terms) of these genes were searched using gseGO (pvalueCutoff=0.05) of clusterProfiler [31]. The resultant top GO terms after simplification (cutoff = 0.6) are shown in Figure 5. The top GO terms are grouped by the distinct ontology terms Biological Process (BP), Cellular Component (CC), and Molecular Function (MF). The full list of genes under each enrichment term is under supplementary Table 1. We found several M band proteins enriched in the muscle cell differentiation, muscle system process, and actomyosin structure organization terms. The muscle cell differentiation term includes processes related to myogenesis that an unspecialized cell would require to differentiate into a muscle cell. Actomyosin structure organization includes processes that are involved in assembly and organization of structures containing actin and myosin. Under these terms, we found *Lmod2* and *Myom3* up-regulated in AD muscles. *Myom1* and *Myom2* are the two prominent isoforms and show no significant differential expression between the AD and SM. Although the absolute expression of *Myom3* is lower than that of *Myom1* and *Myom2* in both tissues, it is significantly upregulated in AD (log2FC = 3.46, padj=5.68E-05). *Lmod2* is a protein that is involved in actin nucleation, thin filament length regulation, and regulation of contractility (log2FC = 1.37, padj = 8.76E-03) [16]. A known co-expressed protein with *Lmod2* is Tropomodulin 1 (*Tmod1*), and it is contrastingly up-regulated in SM muscles (log2FC = 1.02, padj = 2.60E-02). Like *Lmod2, Tmod1* is involved in actin filament length regulation by capping the pointed ends of the thin filament [9]. The thin filament length is dynamically altered due to frequent breakages and regeneration. The SM muscles also had an increased expression of myosin binding protein-C 2 (*Mybpc2*, log2FC = 1.05, padj = 3.30E-02)

**Figure 5.**
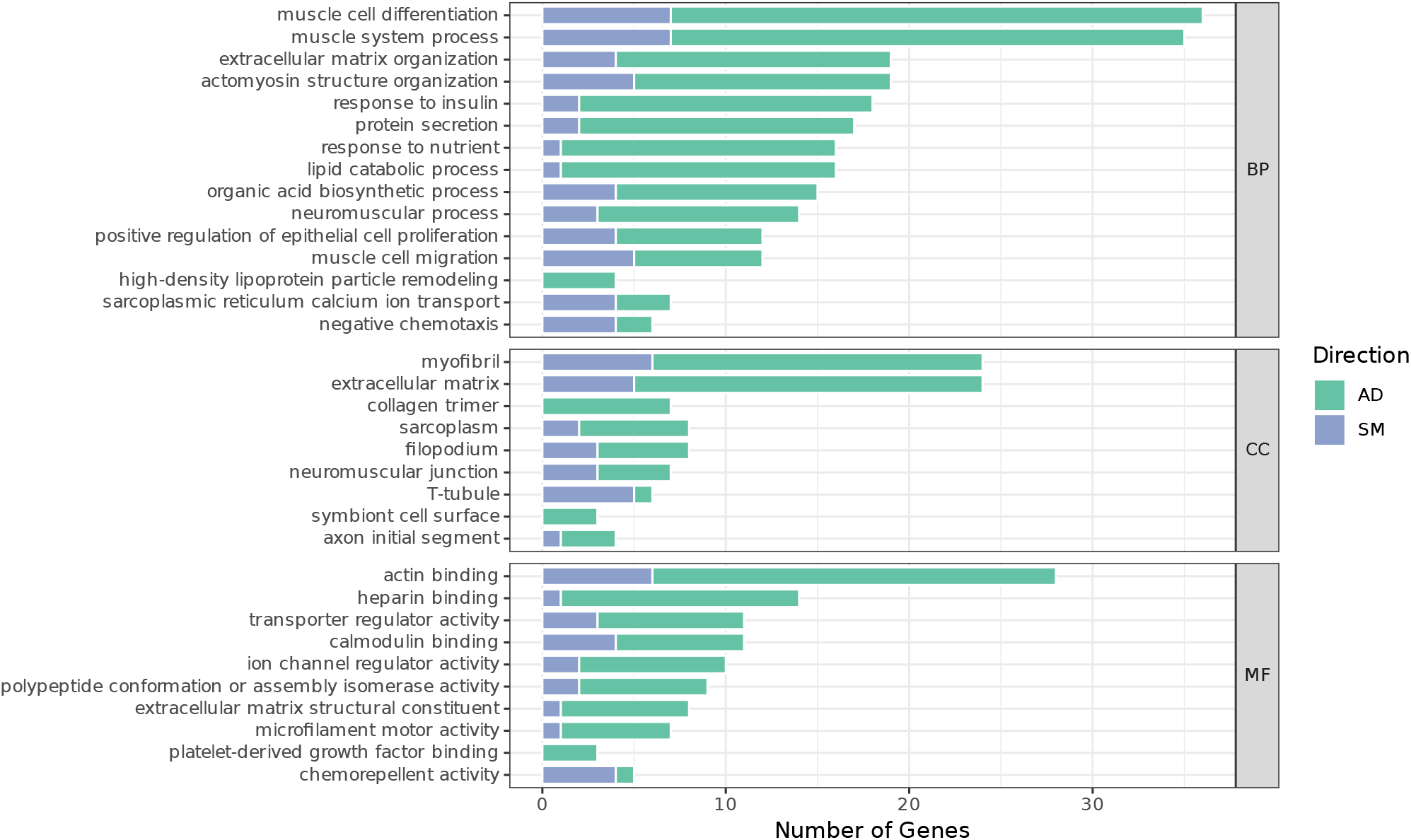
Gene ontology enrichment exhibits an asymmetrical structural terms dominated by jaw opener upregulation in the differentially expressed geneset between SM and AD. GO terms are divided into major non-overlapping gene ontology categories: Biological process (BP), Cellular component (CC) and, Molecular function (MF). Number of differentially expressed genes enriched under each term is shown on the y-axis and the direction of up regulation is colored based on if it is upregulated in AD (green) and SM (blue). Genes upregulated in AD muscles include GO terms related to muscle cell differentiations, muscle system process, actin binding, and calmodulin binding.

An interesting GO term under BP is the ‘sarcoplasmic reticulum calcium ion transport’ which has similar number of genes up regulated in both AD and SM. AD, which reportedly has a higher sensitivity to calcium [23], shows an upregulation in Small Transmembrane Regulator of Ion Transport 1 (*Strit1*), Sarcolipin (*Sln*), and *Atp2a2* genes. *Atp2a2* gene codes for one of the SERCA/Ca2+ ATPases, which is an important enzyme involved in transportation of Ca2+ ions into the sarcoplasmic reticulum from the cytosol. This is known to be the most abundant protein found in the sarcoplasmic reticulum [30]. The *Atp2a2* isoform is the slow twitch type isoform. The SERCA pump activity is regulated by Sln which inhibits the SERCA activity by reducing the maximum calcium uptake rate without affecting the ATP hydrolysis rate. It is reported that muscles over expressing sarcolipin fatigue significantly less and could enhance fatty acid metabolism [28]. *Sln* decoupling the Ca2+ from SERCA leads to increased cytosolic Ca2+ concentration, which in turn has shown to increase mitochondrial metabolism. A major Ca2+ sensitive pathway in skeletal muscles is the Ca2+/calmodulin dependent kinase pathways. Calmodulin dependent protein kinases play a key role in regulating several pathways related to metabolism, signaling, and ion channel regulation. Out of the three calmodulin isoforms, Calmodulin 1 (*Calm1*) is significantly upregulated in SH compared to both AD and SM, and it is lowest in AD (Figure 6b). There were several genes enriched under the calmodulin binding GO term. Wolframin ER Trans-membrane Glycoprotein (*Wfs1*) gene under this GO term produces wolframin which is an endoplasmic reticulum protein that is reported to be indirectly involved in Ca2+ homeostasis and Ca2+ transport between mitochondria and ER. *Wfs1* was significantly up-regulated in AD muscles compared to SM.

**Figure 6.**
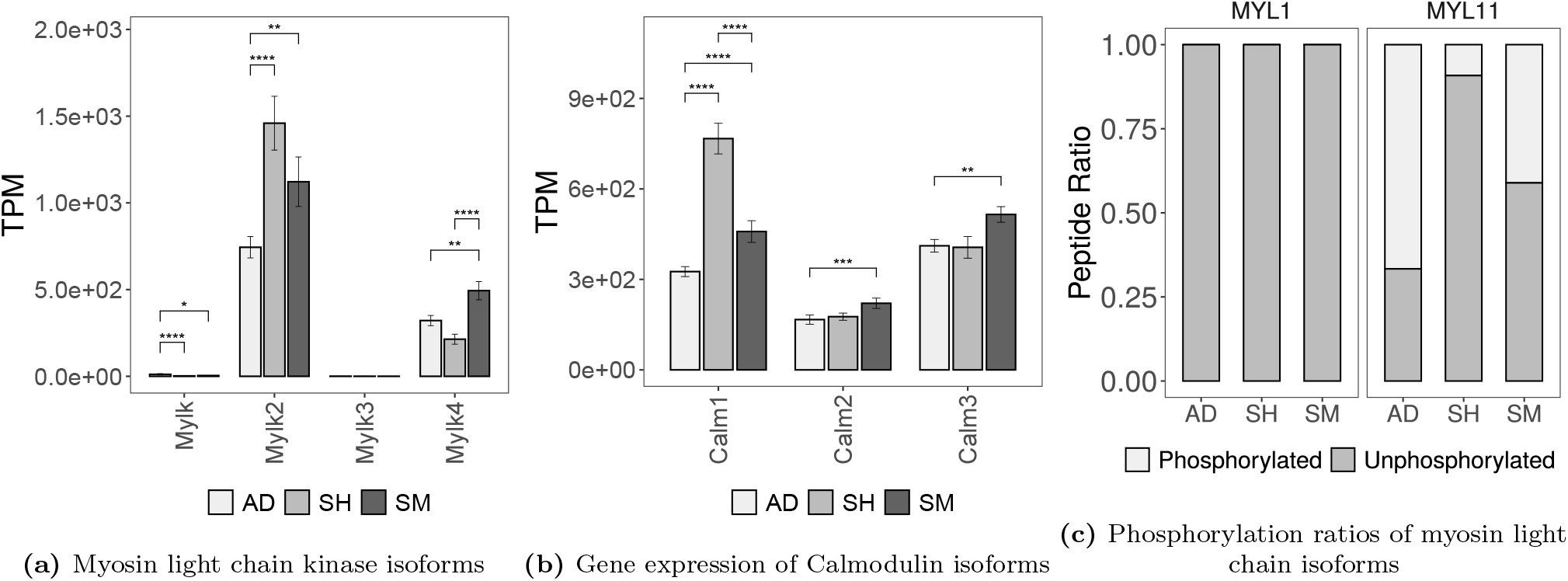
Expression of calcium binding proteins and phosphorylation ratios of MLC in rat jaw muscles. (a) Transcript abundance of *Mylk2* is highest in SH, followed by SM and is lowest in AD. (b) Calmodulin 1 expression follows the same trend observed in *Mylk2*. (c) Phosphorylation ratios of the regulatory light chain MYL11 are inversely proportional to the upstream *Mylk2* and *Calm1* transcript levels. This mismatch suggests that myosin regulatory light chain phosphorylation is not directly correlated to steady state kinase expression or activator transcription.

Ca2+/calmodulin dependent pathways are also involved in myosin light chain phosphorylation, which is known to increase ATPase activity and rate of force production. Myosin light chain phosphorylation is carried out enzymatically by myosin light chain kinases. Myosin light chain kinases are activated by calmodulin, which is activated by binding to Ca2+. Expression of the two major myosin light chain kinases is shown in Figure 6a. Figure 3. 6b shows the calmodulin expression in the three muscles. The expression of *Mylk2* is the highest in SH, followed by SM and is the least in AD. An identical pattern is shown in the expression of *Calm1* in Figure 6b, with SH expression being highest and AD expression being lowest. Figure 6c shows the phosphorylation ratios in Myosin light chain 1 (*Myl1*) and Myosin light chain 11 (*Myl11*). The essential light chain *Myl1* exists fully un-phosphorylated while the regulatory myosin light chain *Myl11* shows variable phosphorylation. The ratio of phosphorylation is the lowest in SH, followed by SM and is the highest in AD. RLC phosphorylation increases myosin duty cycle, which enhances force development while decreasing contractile velocity [11]. As previously discussed, the rat AD exhibits efficient, low intensity, and high frequency contractions, which benefits from this increased duty cycle because it allows the muscle to hold tension with minimized ATP breakdown. However, this phosphorylation pattern of *Myl11* in the three muscles is inverted relative to the upstream *Mylk2* and *Calm1* transcript abundance. This could be a result of a divergence between steady state transcript levels and downstream protein modification pathways. Given the sample size constraints of this exploratory proteomic analysis, expanded validation across a larger sample size remains essential to fully elucidate this post-translational regulatory pathway.

### Somitogenesis related transcription factors are differentially expressed in sternohyoid muscles

The AD and SM muscles share a common embryonic lineage, developing from the first branchial arch whereas the sternohyoid (SH) does not. In addition to that, functionally the SH does not directly participate in mastication. These shared developmental and functional features are reflected in the differential expression analysis; when evaluated against the non-masticatory SH baseline, the AD and SM exhibit a shared DE geneset of 179. The top gene ontology terms (GO terms) enriched within this intersection are shown in Figure 7 with the full list of constituent genes detailed in Supplementary Table 2. The functional enrichment highlights embryonic patterning pathways such as ‘anterior/posterior pattern specification’ and ‘somitogenesis.’ The ‘anterior/posterior pattern specification’ term includes genes that are involved in regionalization processes that differentiate cells along the anterior/posterior axis of an embryo. ‘Somitogenesis’ is a sub GO term ‘under anterior/posterior pattern specification’ and is involved in the formation of mesodermal clusters. Two of the genes under these GO terms are *Dsx* and *Dmrt2* and *Pax3*. The *Pax3* gene is known to be up-regulated in myogenic cell lineages in embryos that target several processes involved in myogenesis, and Dmrt2 is a known target of *Pax3* [26]. Our adult rat SH muscles show a significant down regulation of Dmrt2 and a significant up regulation in *Pax3* compared to both AD and SM. We also observed a significant down regulation of *Six2* in SH which is reported to increase In vitro muscle cell proliferation in bovine skeletal muscle cells. Collectively, this persistent embryonic gene expression retained within adult musculature indicates that these developmental regulators continue to dictate regional identity and patterning across mature tissues of distinct embryological origins.

**Figure 7.**
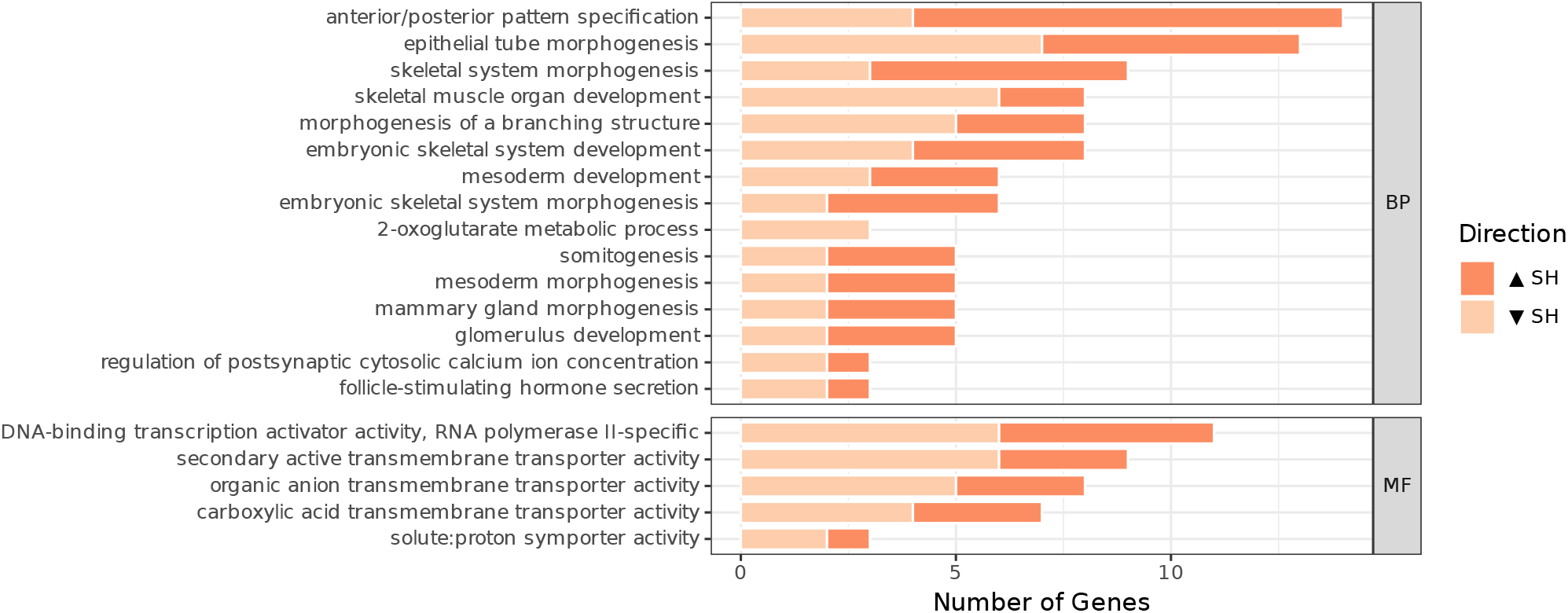
Enriched biological process and molecular function categories derived from the intersection of 179 common differentially expressed genes identified in pairwise comparison of first branchial arch muscles against the SH. The enriched terms demonstrate that adult masticatory muscles retain persistent transcriptomic expression of their branchial specification genes.

### Network analysis shows AD muscles show elevated mitochondrial, vascular, and lipid metabolism genes

Network of co-expressed genes created using Weighted Gene Co-expression Analysis (WGCNA) involves calculating pairwise correlation coefficients, where stronger connections are amplified by raising the coefficients to a soft thresholding power; this value is known as the co-expression weight and is elevated in highly co-expressed genes. The strengths of these weights are used to determine the neighborhood of a gene (topological overlap). A gene module is a highly interconnected cluster of genes with an elevated level of co-expression. Because the genes in a gene module are more likely to be involved in a shared biological pathway. Each co-expression module is characterized by module eigengenes that represents its collective expression pattern. These eigengene profiles are compared across the samples to calculate the correlation between the modules and the tissue types. The WGCNA analysis resulted in 17 modules and Figure 8 shows the relationship between the module eigengenes, and the three muscle tissue types and sex. The values inside the boxes indicate the correlation coefficient, and the number within the brackets shows the significance. AD positive modules are darkolivegreen (r = 0.69, p = 5 × 104), brown (r = 0.61, p = 0.003), cyan (r = 0.63, p = 0.002) and paleturquoise (r = 0.95, p = 7 × 1011). Floralwhite is positively correlated in SH (r = 0.92, p = 3 × 109) and the modules darkolivegreen (r = 0.59, p = 0.005), brown (r = 0.81, p = 8 × 106) and lightyellow (r = 0.82, p = 5 × 106) are SH negative. The SM positive module is lightyellow (r = 0.81, p = 7 × 106) and cyan (r = 0.95, p = 6 × 1011) is SM negative. Module associations with sex as a trait were non-significant in all recognized modules, suggesting that the transcriptomic architecture of these jaw muscles is driven by tissue type and function rather than sexual variance.

**Figure 8.**
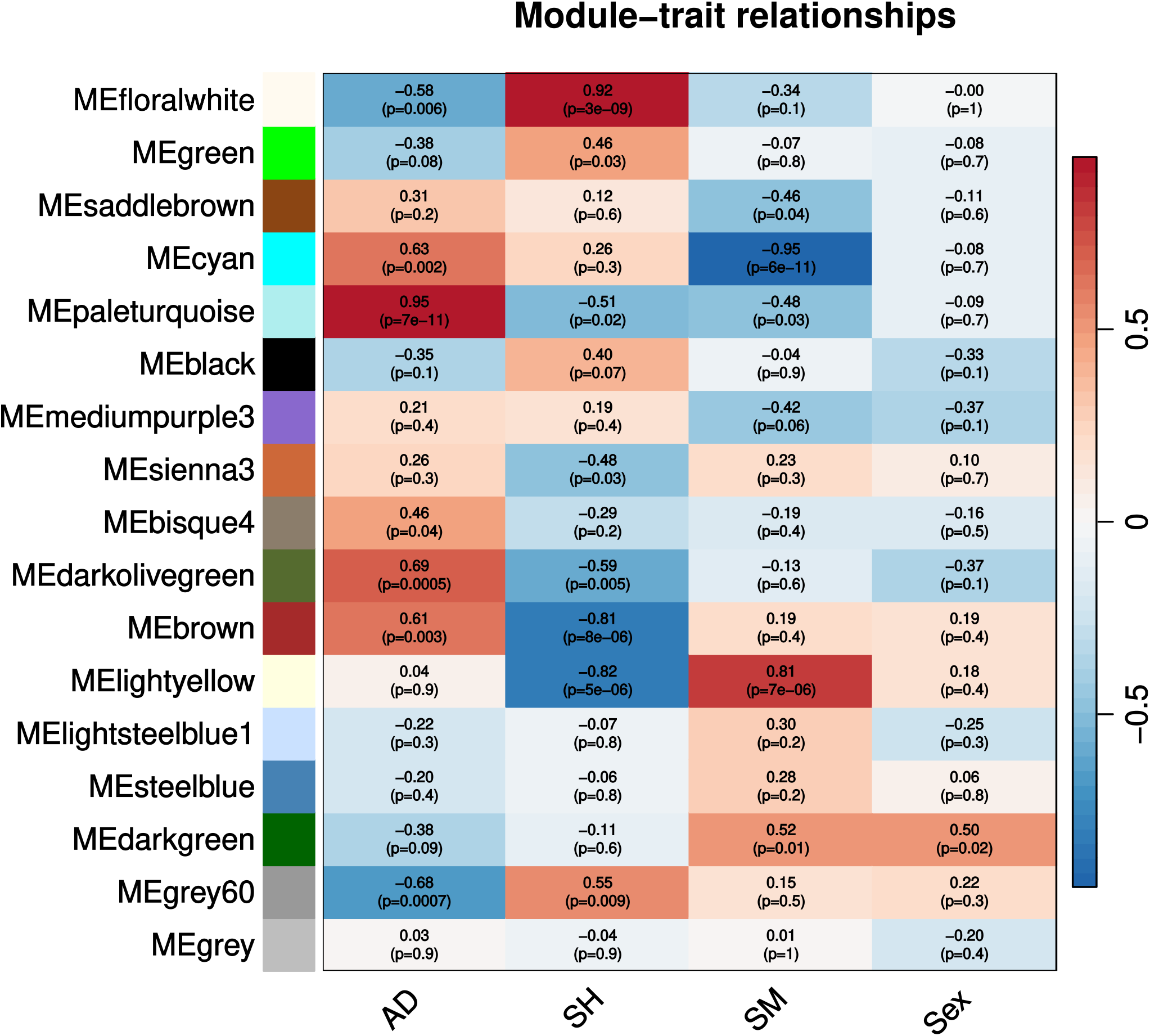
Heatmap of module and trait correlations in rat jaw muscle demonstrates that the paleturquoise, brown, and cyan modules are highly correlated with the AD, SH, and SM samples respectively. Each row represents a co-expression module (labeled by color), and each column represents the sample traits (AD, SH, SM, and Sex). Numbers within the cells indicate the Pearson correlation coefficient, with the respective p-values in parentheses. The color scale indicates the strength and the direction of the correlation: red represents a positive correlation (upregulation of module genes in samples with the trait), and blue represents a negative correlation (downregulation of module genes with respect to the trait). The cyan module shows a strong negative correlation with the SM (*r* = −0.95, *p* = 6 *×* 10^−11^) and paleturquoise module is strongly positively correlated with the AD (*r* = 0.95, *p* = 7 *×* 10^−11^).

The GO terms of each module are shown in Figures 9 and 10. The brown module, which was strongly negatively correlated with SH samples, was enriched for GO terms associated with mitochondrial and oxidative metabolism. The top biological process (BP) GO terms in the brown module contained cellular respiration, mitochondrial respiratory chain complex assembly, mitochondrial transport, and mitochondrial gene expression and the cellular component (CC) GO terms included mitochondrial protein-containing complex. Molecular function (MF) GO terms included electron transfer activity and, oxidoreduction-driven active trans-membrane transporter activity. In summary, these enriched terms suggest that genes involved in mitochondrial energy production are down-regulated in SH relative to AD.

**Figure 9.**
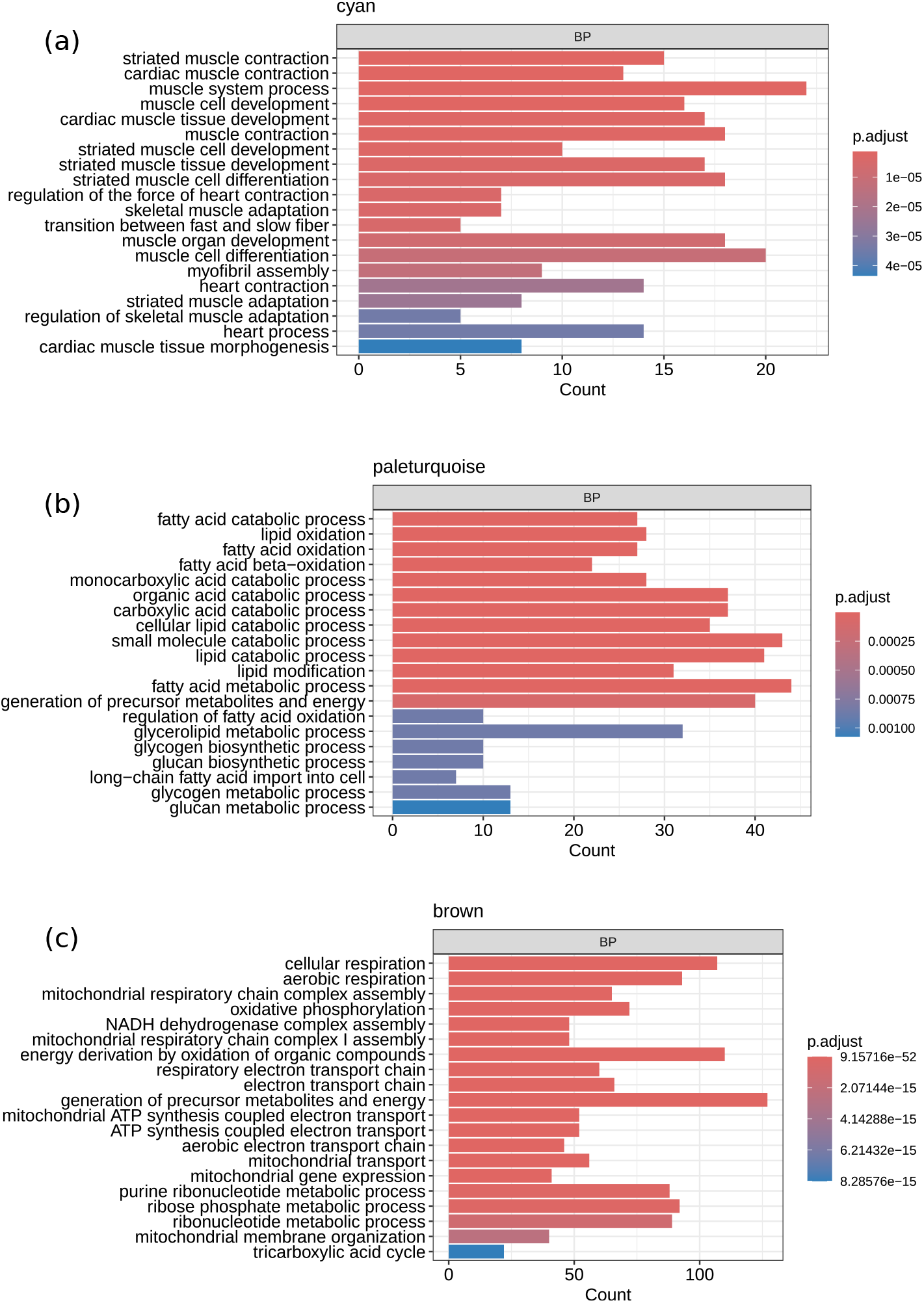
Functional enrichment of WGCNA modules associated with AD, SH and SM muscles of the rat. The cyan module was positively correlated with AD and negatively correlated with SM. Enriched ontology terms were related to muscle contraction, development, and differentiation. (b) The paleturquoise module was also positively correlated with AD and negatively correlated with SM. This module was enriched for pathways associated with fatty acid and glycogen metabolism, and insulin response. (c) The brown module was positively correlated with AD and negatively correlated with SH muscles. Genes within this module were enriched in cellular respiration, aerobic respiration, mitochondrial organization, and transport related processes. The color scale represents the statistical significance (adjusted p-value) of each ontology term and the bar length represents the number of genes enriched within each term.

**Figure 10.**
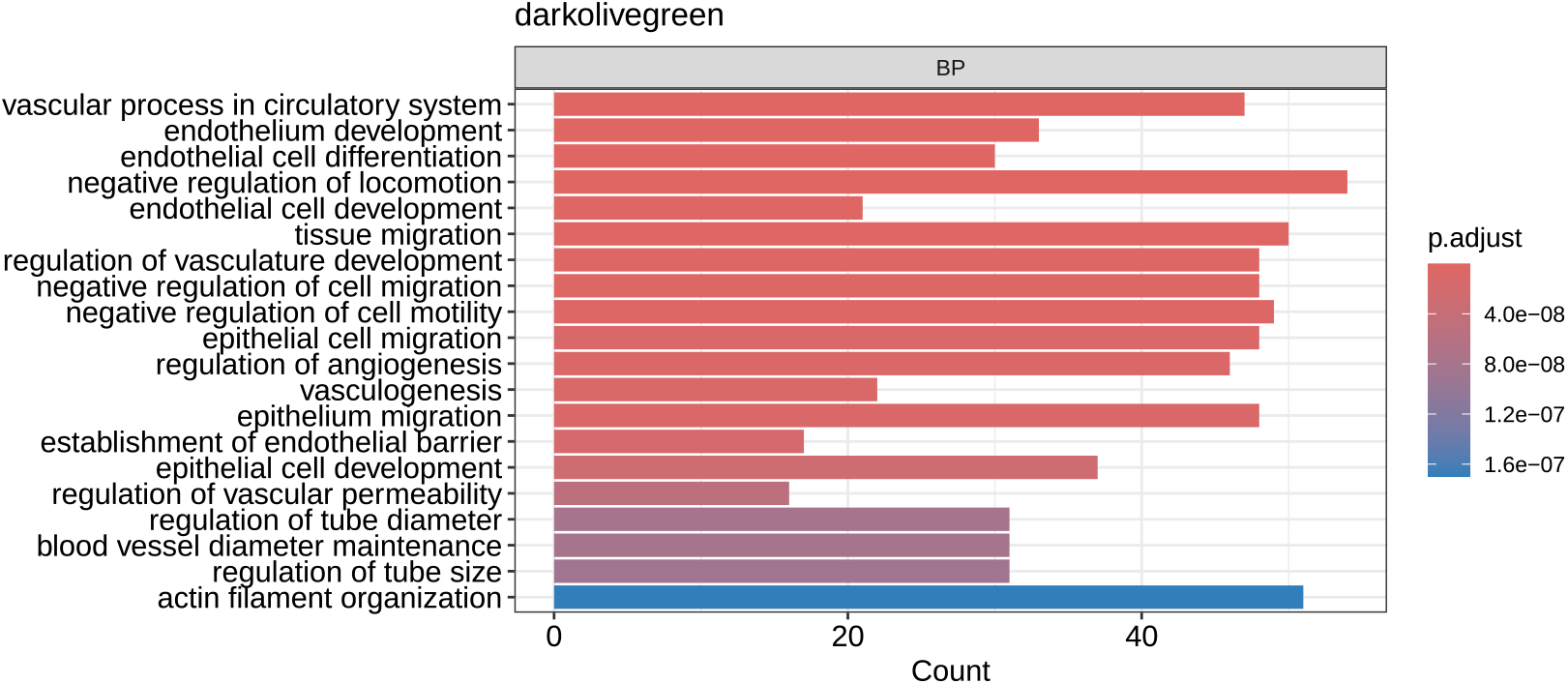
Gene ontology enrichment of the darkolivegreen co-expression module highlighting vascular and structural divergence between AD and SH. Characterized by a positive correlation with the AD and a negative correlation with the SH, this module displays enrichment for vascular and extractellular matrix related processes. These enriched categories demonstrate that AD tissues are characterized by an upregulation of genes dedicated to ECM remodeling and vascular network organization, further highlighting the aerobic nature of AD muscles, which require an extensive vascular network to secure sustained oxygen supply. Bars indicate the gene count included within each ontology term, and the color scale indicates the statistical significance (adjusted p-value)

The cyan module is positively correlated with AD and negatively correlated with SM, reflecting that some of the contrasting variability between a jaw opener and a jaw closer muscle. Gene enrichment of the cyan module genes included striated muscle contraction, muscle system process, and muscle cell development as the top GO terms in biological process and in the cellular component related GO terms contractile fiber and myosin complex. They also show actin binding and cytoskeletal motor activity related GO terms in molecular Function.

Paleturquoise is positively correlated with AD and negatively correlated with both SH and SM. The paleturquoise module genes are enriched with biological process GO terms related to metabolism of fatty acids, glycogen, glucan, carnitine, purine ribonucleotides, alcohol, and response to insulin. Cellular component GO terms contain nucleoid and mitochondrial nucleoid. Molecular function GO terms are oxidoreductase activity, calmodulin-dependent protein kinase activity, and fatty acid derivative binding. Notable genes in fatty acid catabolic process pathways that were up-regulated in AD included Acetyl-CoA Acyl transferases such as Acaa2 and Acyl-CoA Dehydrogenase genes such as *Acadl, Acadm* and *Acadvl*. Hydroxyacyl-CoA Dehydrogenases such as *Hadha*and *Hadhb*were also significantly up-regulated in AD. The *Acadl, Acadvl* and *Acadm*also perform a response to cold function.

The darkolivegreen module, positively correlated with AD, displayed enrichment for vascular and extracellular-matrix (ECM) related processes. The top BP-GO terms included the vascular process in the circulatory system, endothelium development, and negative regulation of locomotion. CC-GO terms included actin filament bundle and receptor complex. MF-GO terms contain cell adhesion molecule binding, trans-membrane receptor protein kinase activity, and GTPase regulator activity and GTP binding. These terms imply that AD samples exhibit an up-regulation of ECM and vascular organization-related genes.

According to these results, the AD exhibits a distinct expression profile characterized by specialized energy substrate utilization and elevated mitochondrial energy production. Specifically, the AD shows elevated expression of genes enriching both lipid and glycogen metabolism. Glycogen provides access to rapid energy production, and fatty acid metabolism allows flexibility in available energy resources.

## Discussion

Despite functional and developmental differences, all three tissues utilized in this study exhibited a principally fast contractile profile; characterized by the high expression of the canonically fast-twitch isoforms *Myh1, Myh4, Myl1*, and *Myl11*. Functional divergence between these tissue types is likely driven by variations in overall composition. The jaw-opening AD showed elevated expression of slow and oxidative contractile isoforms including the slow-type I *Myh7* and oxidative-glycolytic *Myh2* and slow troponin isoforms. Together, these fatigue resistant isoforms accounted for 13% of the total myosin heavy chain isoforms in the AD, compared to below 5% in the other muscles. *Myh7* represented only 2% of the total myosin heavy chain transcript expression in the AD, whereas a previous protein level study reported 10% MHC-I in the rat AD [23]. This discrepancy could be related to translational efficiency, protein turnover rates as well as differences in rat strain, age or diet between the studies. However, the greater expression of slow and oxidative isoforms in the AD is consistent with the repetitive, sustained contractions required of the rat AD in the stabilization and postural maintenance of the mandible [5, 23]. In contrast, the rat SM had the least amount of slow *Myh7* expression and displayed a predominantly fast glycolytic fiber type optimized for phasic, high force tasks associated with biting down on hard rodent chow. This isoform distribution is different from that reported in human jaw muscles, in which jaw-closing muscles contain a greater proportion of slow MHC-I fibers, and jaw-opening muscles predominantly express fast-type MHC-II fibers [13]. This interspecies difference has been attributed to the role of the human jaw-closing musculature in stabilizing the mandible and maintaining its posture against gravity [23]. This elevated expression of slow contractile isoforms also extends to the proteins in the troponin complex. Fast troponin isoforms were dominant in all three muscles, however the AD exhibited a relative higher expression of slow isoforms *Tnnc1, Tnni1*, and *Tnnt1*.

The specialization of the AD extends beyond contractile isoform usage; our network analysis reveals that these sarcomeric adaptations in the AD are supported by a coordinated upregulation of metabolic and structural infrastructures. Network modules highly positively correlated with the AD (paleturquoise, darkolivegreen, and brown modules in Figure 8) are enriched for genes governing oxidative metabolism, angiogenesis, and the metabolic processing of a variety of substrates. The paleturquoise module has the highest positive correlation with AD (r = 0.95) and is enriched in fatty acid catabolic enzymes (*Acaa2, Acadm, Acadl, Acadvl, Hadha*, and *Hadhb*). This indicates that the AD possesses the ability to utilize both glycogen and fatty acids as substrates for energy production, aligning with its higher expression of aerobic myosin isoforms. These paleturquoise module genes are significantly suppressed in the SH and SM samples, which show minimal expression of aerobic myosin isoforms. As discussed previously, major fast-twitch isoforms of myosins, myosin light chains, tropomyosins and troponins are significantly downregulated in the AD. This reduced expression of major contractile proteins may reflect the specialization of the AD for efficient, sustained, low-intensity activity and its correspondingly higher duty time. Although not directly analogous to present transcriptomic findings, evidence from both human and rodent models indicate that prolonged muscle activity is associated with reduced global protein turnover [25]. Contractile specialization may also involve differences in calcium responsiveness. During calcium-activate contraction, Ca^2+^ ions bind to troponin C, causing a conformational change in the troponin-tropomyosin complex, composed of troponin C, troponin I and troponin T. Troponin T binds to tropomyosin and is often correlated with calcium sensitivity [29]. The elevated expression of slow troponin isoforms in the AD may contribute to its higher calcium sensitivity and support sustained contraction at relatively low activation levels. However, the cyan module is positively correlated with the AD (r = 0.63) and represents developmental genes promoting myofibril assembly, muscle cell development, and contraction. Specifically, this module comprises slow-twitch variants and minor contractile proteins rather than the predominant fast-twitch isoforms that characterize the architecture of these tissues. In contrast, this module is significantly negatively correlated with the SM (r = −0.95), suggesting a suppression of these enriched pathways. Additionally, the lightyellow module, which is significantly positively correlated with SM, lacks any significantly enriched functional terms. The combination of this lack of functional enrichment among upregulated genes in SM and the suppression of muscle developmental genes suggests that SM does not need to continuously express a diverse set of contractile transcripts. This transcriptomic profile aligns with previous physiological studies characterizing rat SM contractions as short, rapid bursts. Such contractions require less slow-twitch isoform expression compared to the chronically active, stabilizing mechanics of the AD.

Like SM, the SH maintains a low isoform diversity in its expression. Its myosin isoform composition includes 79% of just *Myh4*. It is also significantly negatively correlated in metabolic processes related paleturquoise module, mitochondria and aerobic respiration related brown module. This demonstrates that the angiogenesis, aerobic respiration and metabolism related traits observed in the AD are not characteristic of the SH. SH exhibits a higher overall differential expression compared to other two muscles, and 179 genes were commonly differentially expressed in both the AD and SM when compared to the SH (Figure 4). These comparisons suggest that the common branchial arch origin of AD and SM might be driving this divergence. The functional enrichment of these 175 shared genes (Figure 7) highlighted development specific pathways named ‘anterior/posterior pattern specification’ and ‘somitogenesis’. Within these pathways we identified an upregulation of *Pax3* and a downregulation of its known downstream target *Dmrt2*. These genes determine regionalization during embryonic stages, and their persistent expression in the adult rat was surprising, suggesting that regionalization-related genes are still maintained in the adult.

In conclusion, our transcriptomic analyses reveal that the specialization of jaw musculature is not limited to fiber-type classifications. The AD, SH, and SM all maintain a general fast-twitch isoform expression; however, their functional divergence is achieved though fine-tuned shifts in contractile isoform expression and metabolic processes. The SM is geared toward a glycolytic architecture optimized for rapid short bursts of bite force, while the jaw opener is adapted to be metabolically flexible and efficient. Together, these transcriptomic landscapes highlight the antagonistic biomechanics of the feeding apparatus.

