## Supplementary material for "Transcriptomic profile of a rat jaw opener (anterior digastric) and a jaw closer (superficial masseter)": Table 1 and 2

### Supporting Information

**Table 1.** GO terms of Rat SM vs. AD comparison

| GO:ID | Term | geneID | Ratio | p.adj | N | Ont |
|---|---|---|---|---|---|---|
| 0042692 | muscle cell differentiation | Hey2, Tnnt1, Mybpc2, Csrp3, Pak1, Igf2, Prkg1, Ankrd2, Myoz2, Camk2d, Pitx2, Actc1, Lmod2, Tmod1, Myom3, Cfd, Barx2, Cdon, Rbpms2, Myh10, Tbx1, Atp2a2, Myl2, Tbx3, Mybph, Casq1, Meis1, Myh6, Myh7, Fgf9, Comp, Adprhl1, Ninj1, Irx3, Agt, Pi16 | 36/291 | 0.000 | 36 | BP |
| 0003012 | muscle system process | Hey2, Tnnt1, Mybpc2, Csrp3, Pak1, Prkg1, Strit1, Tpm3, Myoz2, Camk2d, Scn1a, Actc1, Myh7b, Lmod2, Lmcd1, Tmod1, Myl6bl1, Sln, Myl3, Chrnd, Myh8, Atp2a2, Myl2, Tbx3, Tnni1, Casq1, Meis1, Myh6, Myh7, Atp8a2, Tnnc1, Comp, Smpd3, Agt, Pi16 | 35/291 | 0.000 | 35 | BP |
| 0031032 | actomyosin structure organization | Tnnt1, Mybpc2, Csrp3, Pak1, Pdlim1, Myoz2, Actc1, Ppp1r9a, Lmod2, Tmod1, Myom3, Myh10, Ppm1e, Myl2, Mybph, Casq1, Myh6, Myh7, Adprhl1 | 19/291 | 0.000 | 19 | BP |
| 0030198 | extracellular matrix organization | Postn, Col11a1, Col1a2, Adamtsl5, Fbln1, Adamts20, Thsd4, Col3a1, Col1a1, Ccdc80, Lamb3, Comp, Mkx, Gfod2, Smpd3, Agt, Col18a1, Spock2, Gpm6b | 19/291 | 0.000 | 19 | BP |
| 0014812 | muscle cell migration | Pak1, Prkg1, Postn, Camk2d, Itga4, Myo5a, Pax3, Adipoq, Slit2, Fgf9, Net1, Agt | 12/291 | 0.000 | 12 | BP |
| 0070296 | sarcoplasmic reticulum calcium ion transport | Strit1, Camk2d, Sln, Mettl21c, Cacng1, Atp2a2, Casq1 | 7/291 | 0.000 | 7 | BP |
| 0050905 | neuromuscular process | Tnnt1, Strit1, Scn1a, Map1a, Myo5a, Chrnd, Myh8, Myh10, Tnni1, Casq1, Myh7, Atp8a2, Tnnc1, Comp | 14/291 | 0.000 | 14 | BP |
| 0032868 | response to insulin | Apoe, Csrp3, Pak1, Ucp2, Eef2k, Igf2, Rbp4, Hmgcs2, Prkcz, Myo5a, Ptprn, Col1a1, Socs3, Bdh1, Adipoq, Slc27a1, Lpl, Agt | 18/291 | 0.000 | 18 | BP |
| 0016042 | lipid catabolic process | Apoe, Hsd3b7, Oxct1, Nceh1, Cidec, Acer2, Endou, Pla2g7, Adipoq, Ehhadh, Scarb1, Acox2, Cyp4f18, Lpl, Lipg, Smpd3 | 16/291 | 0.001 | 16 | BP |

| | | | | | | |
|---|---|---|---|---|---|---|
| 0007584 | response to nutrient | Oxct1, Postn, Lrat, Hmgcs2, Pitx2, Tspo, Cdon, Aldh1a2, Aldh3a1, Col1a1, Bdh1, Adipoq, Scarb1, Lpl, Lipg, C2 | 16/291 | 0.001 | 16 | BP |
| 0050919 | negative chemotaxis | Sema3d, Epha7, Sema7a, Sema6b, Lgr6, Slit2 | 6/291 | 0.002 | 6 | BP |
| 0030016 | myofibril | Tnnt1, Mybpc2, Csrp3, Pak1, Pdlim1, Ankrd2, Myoz2, Scn1a, Actc1, Myh7b, Lmod2, Tmod1, Myom3, Twf2, Myl3, Myh8, Myl2, Mybph, Tnni1, Casq1, Myh6, Myh7, Tnnc1, Adprhl1 | 24/302 | 0.000 | 24 | CC |
| 0031012 | extracellular matrix | Thbs2, Apoe, Otog, Postn, Col11a1, Hmcn2, Angpt4, Col1a2, Mmrn1, Adamtsl5, Fbln1, Adamts20, Thsd4, Col3a1, Col1a1, Ccdc80, P3h2, Lamb3, Slit2, Fgf9, Comp, Fgf1, Gfod2, Col18a1 | 24/302 | 0.000 | 24 | CC |
| 0005581 | collagen trimer | Col11a1, Col1a2, Col3a1, Col1a1, Adipoq, C1qtnf7, Col18a1 | 7/302 | 0.001 | 7 | CC |
| 0016528 | sarcoplasm | Ankrd2, Strit1, Camk2d, Sln, Myh10, P3h2, Atp2a2, Casq1 | 8/302 | 0.001 | 8 | CC |
| 0030175 | filopodium | Cd302, Actc1, Ppp1r9a, Myo5a, Twf2, Fgd4, Spata13, Ninj1 | 8/302 | 0.002 | 8 | CC |
| 0030315 | T-tubule | Camk2d, Kcnj3, Scn1a, Cacng1, Casq1, Stbd1 | 6/302 | 0.006 | 6 | CC |
| 0031594 | neuromuscular junction | Postn, Camk2d, Ppp1r9a, Epha7, Cib2, Chrnd, Myh10 | 7/302 | 0.006 | 7 | CC |
| 0043194 | axon initial segment | Lgi1, Camk2d, Scn1a, Map1a | 4/302 | 0.006 | 4 | CC |
| 0106139 | symbiont cell surface | C2, C4a, C4b | 3/302 | 0.021 | 3 | CC |
| 0003779 | actin binding | Mybpc2, Csrp3, Mical2, Pdlim1, Ablim1, Tpm3, Ankrd35, Myoz2, Map1a, Myh7b, Ppp1r9a, Lmod2, Tmod1, Inf2, Panx1, Myo5a, Twf2, Myl3, Myh8, Myh10, Fgd4, Myl2, Tnni1, Vash2, Afap1, Myh6, Myh7, Tnnc1 | 28/290 | 0.000 | 28 | MF |
| 0008201 | heparin binding | Rspo3, Thbs2, Apoe, Postn, Col11a1, Adamtsl5, Ccdc80, Lgr6, Slit2, Fgf9, Comp, Lpl, Fgf1, Lipg | 14/290 | 0.000 | 14 | MF |
| 0000146 | microfilament motor activity | Actc1, Myh7b, Myo5a, Myh8, Myh10, Myh6, Myh7 | 7/290 | 0.000 | 7 | MF |
| 0005201 | extracellular matrix structural constituent | Otog, Col11a1, Col1a2, Fbln1, Col3a1, Col1a1, Comp, Col18a1 | 8/290 | 0.001 | 8 | MF |
| 0120544 | polypeptide conformation or assembly isomerase activity | Dnah5, Actc1, Myh7b, Kifc2, Myo5a, Myh8, Myh10, Myh6, Myh7 | 9/290 | 0.005 | 9 | MF |
| 0045499 | chemorepellent activity | Sema3d, Epha7, Sema7a, Sema6b, Slit2 | 5/290 | 0.005 | 5 | MF |

| | | | | | | |
|---|---|---|---|---|---|---|
| 0141108 | transporter regulator activity | Prkg1, Kcnab1, Camk2d, Prkcz, Fxyd6, Lrg1, Atp1b2, Cacng1, Atp2a2, Kcng4, Agt | 11/290 | 0.012 | 11 | MF |
| 0005516 | calmodulin binding | Eef2k, Syt7, Camk2d, Myh7b, Myo5a, Myh10, Wfs1, Myh6, Myh7, Phkb, Grm4 | 11/290 | 0.014 | 11 | MF |
| 0099106 | ion channel regulator activity | Prkg1, Kcnab1, Camk2d, Prkcz, Fxyd6, Lrg1, Cacng1, Atp2a2, Kcng4, Agt | 10/290 | 0.017 | 10 | MF |
| 0048407 | platelet-derived growth factor binding | Col1a2, Col3a1, Col1a1 | 3/290 | 0.020 | 3 | MF |

**Table 2.** GO terms of non-orthologous genes only found in the rat genome

| | | | | | | |
|---|---|---|---|---|---|---|
| 0097696 | cell surface receptor signaling pathway via STAT | Clcf1, Cd40, Socs2, Naglu, Csf2ra, Il15ra, Tslp, Arl2bp | 8/194 | 0.049 | 8 | BP |
| 0002819 | regulation of adaptive immune response | Raet1ll1, Clcf1, Il1b, Cd40, Mad2l2, Tnfrsf14, Cd7, Muc4, Tnfsf18, Cd24 | 10/194 | 0.049 | 10 | BP |
| 0019730 | antimicrobial humoral response | Wfdc15b, Wfdc2, Ccl27, Ccl25, Cxcl13, Rnase3, Defb1, Tslp | 8/194 | 0.049 | 8 | BP |
| 0042742 | defense response to bacterium | Il1b, Wfdc15b, Wfdc2, Tnfrsf14, Ngp, Ccr5, Cxcl13, Rnase17, Rnase3, Gpr15lg, Defb1, Tslp | 12/194 | 0.049 | 12 | BP |
